# A Comparative Study of Radiomic and Connectomic Approaches to Classification of IDH1 Status and 1p/19q Co-deletion in Lower Grade Gliomas

**DOI:** 10.1101/2024.09.14.613034

**Authors:** Rohit V. Paradkar, Ron L. Alterman

## Abstract

**Purpose:** Grade III and IV brain tumors are labeled “high grade”, or malignant. Lower grade tumors (grade II and III) can progress to high grade and must be closely monitored. In lower grade gliomas, the presence of a specific IDH1 gene mutation and the 1p/19q chromosomal co-deletion confer favorable prognosis and alternative treatment strategy. Presently, these markers are evaluated using surgically obtained tissue specimens. In this study, we evaluate noninvasive approaches to classification of these genetic markers. We hypothesized that connectomic and radiomic approaches to classification would perform similarly. We also tested combined classification, incorporating radiomics and connectomics.

**Methods:** Binary classifiers used radiomic and connectomic features from MRI to classify IDH1 and 1p/19q co-deletion status. Radiomic features were calculated to characterize tumor gray-level, texture, and shape. Voxel-based morphometry was performed to create gray-matter structural connectomes. Nodal efficiencies of brain regions, number of nodes and connections were computed. Binary classifiers predicted IDH1 and 1p/19q co-deletion status. Statistical analysis quantified differences in model performance.

**Results:** Connectomic and radiomic features had insignificant difference in classification of IDH1 status. Radiomic and connectomic classification of 1p/19q co-deletion status had no significant accuracy difference, however, radiomics had significantly higher AUC score. The combined approach had no significant difference to radiomics and connectomics except for a significantly higher AUC score than connectomics in 1p/19q co-deletion classification.

**Conclusion:** Altogether, the study shows that radiomics, connectomics, and a combination of the two are viable classification approaches for these markers. Future studies could incorporate these methods to improve diagnostic performance.

## Introduction

Gliomas are the most common primary tumor of the human brain. In 2016, the World Health Organization (WHO) updated its classification system for gliomas to include genomic and molecular parameters. The WHO system recognizes four histologically distinct grades of glioma (I-IV).^1^ Grade II and III gliomas are termed ‘lower grade gliomas’, or LGGs, and are distinct from Grade IV gliomas, or glioblastomas (GBM), which are among the most malignant and deadly forms of human cancer. While LGGs are not as aggressive as GBM per se, it is important to understand their biology as they have the potential to transform into GBM over time. As one can see in Figure 1, molecular and genomic tumor markers are central to the WHO classification. Per this system, LGGs are associated with a mutation of the isocitrate dehydrogenase 1 (IDH1) gene, which is present in more than three-quarters of these tumors.^3^ The wild type IDH1 gene encodes the α-Kg enzyme, which is a key factor in some metabolic pathways, cell homeostasis, and cellular growth. However, when a missense mutation converts arginine to histidine at residue 132, the IDH1 gene instead encodes the 2-HG enzyme, a known onco-metabolite.^4^ Consequently, IDH1 mutations have the potential to drive gliomagenesis. A second important genetic marker for the LGG subtype is the 1p/19q chromosomal co-deletion. The vast majority of 1p/19q co-deleted tumors are also found to have the IDH1 mutation.^5^ In addition to enabling classification as an LGG, both the 1p/19q co-deletion and IDH1 mutation are important prognostic indicators. Specifically, these genetic markers are associated with improved life expectancy and responses to specific chemotherapeutic agents.^6^ Therefore, testing for these genetic markers is important for determining diagnosis, prognosis, and the appropriate treatment plan for patients suffering with these tumors.

**Figure 1.**
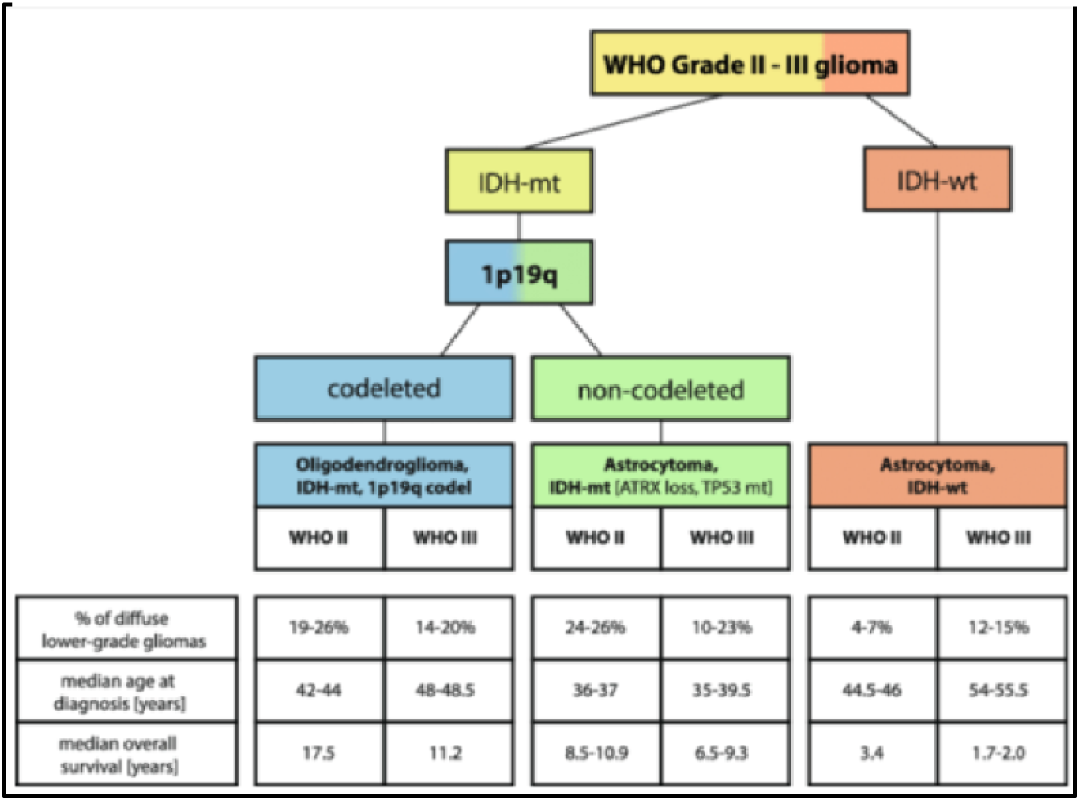
World Health Organization classification of lower grade gliomas based on molecular and genetic parameters^2^.

Current methods of testing for these markers requires invasive brain surgery to obtain tumor tissue.^7^ Genetic testing is expensive, and many hospitals lack the capacity to perform these analyses themselves.^8^ In recent years, several researchers have developed digital imaging methods to classify a tumor’s IDH1 and 1p/19q co-deletion status quickly and non-invasively. One such approach, termed radiomics, utilizes discernible features of the patient’s MRI (Figure 2a). These features include tumor shape, texture, gray-level, and many other features that are imperceptible to the human eye but easily detected through computer-aided applications. Lu et al. employed a radiomics approach to achieve a 91.6% IDH1 classification accuracy and an 87.7% accuracy for 1p/19q classification.^9^ Other studies have achieved similar success, making the radiomics method a promising alternative to invasive, tissue-dependent classification.^10, 11^

**Figure 2.**
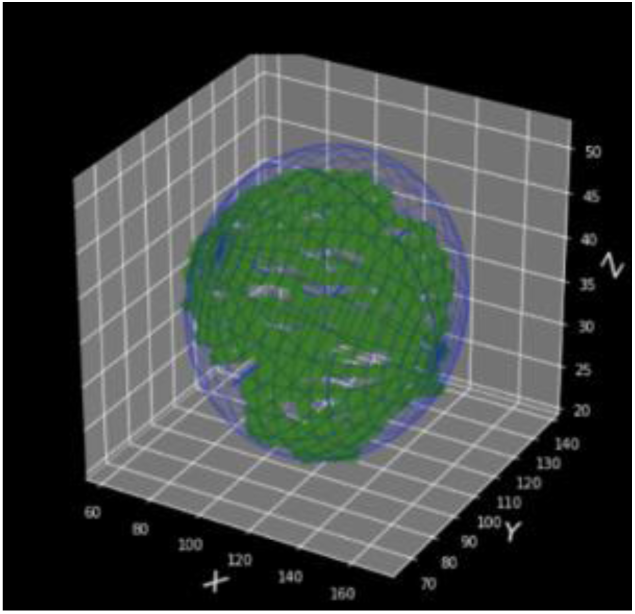
Three-dimensional tumor enclosed in its minimum bounding ellipsoid.

**Figure 2b.**
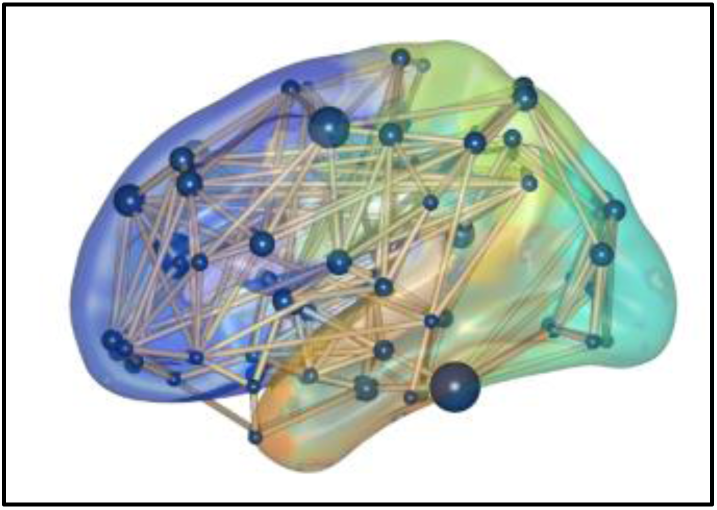
Ball-and-stick representation of a connectome.

Connectomics is a newer and less utilized approach. A connectome is a neural network based on graph theory that models the brain as a collection of ‘nodes’ representing neurons and ‘edges’, which represent the connections between those neurons (Figure 2b). The rationale behind the using a connectomic approach to tumor classification is that focal tumors, such as LGGs, are known to affect entire brain networks. As the connectome is impacted by both biological and epigenetic factors, IDH1 and 1p/19q co-deletion may cause changes to the connectome in a distinct, observable manner. Prior studies support this line of reasoning.^12, 13^ For example, Kesler et. al. employed voxel-based morphometry (VBM) and computed nodal efficiencies to construct connectomes for 90 brain regions using the Automated Anatomical Labeling (AAL) parcellation scheme. Using these and three other connectomic features, their study classified IDH1 status with 86% accuracy.^14^

The aim of this study was to compare radiomic and connectomic approaches to IDH1 and 1p/19q co-deletion classification based on an objective set of criteria and to explore whether the two methods might provide complementary information to enhance diagnostic accuracy.

## Methods

### Overview

The study workflow is depicted in Figure 3. In brief, we extracted radiomic and connectomic features from the same imaging dataset and used those features as input to train and test binary classifiers to predict IDH1 and 1p/19q status. We hypothesized that radiomic and connectomic classification would yield similar results because, although there is a larger existing body of knowledge about the association of radiomic features and prognostic biomarkers like 1p/19q and IDH1, using a connectomic method also seems to be a very usable application considering that focal tumors such as LGG have been found to affect gray-matter covariance networks.^12^

**Figure 3.**
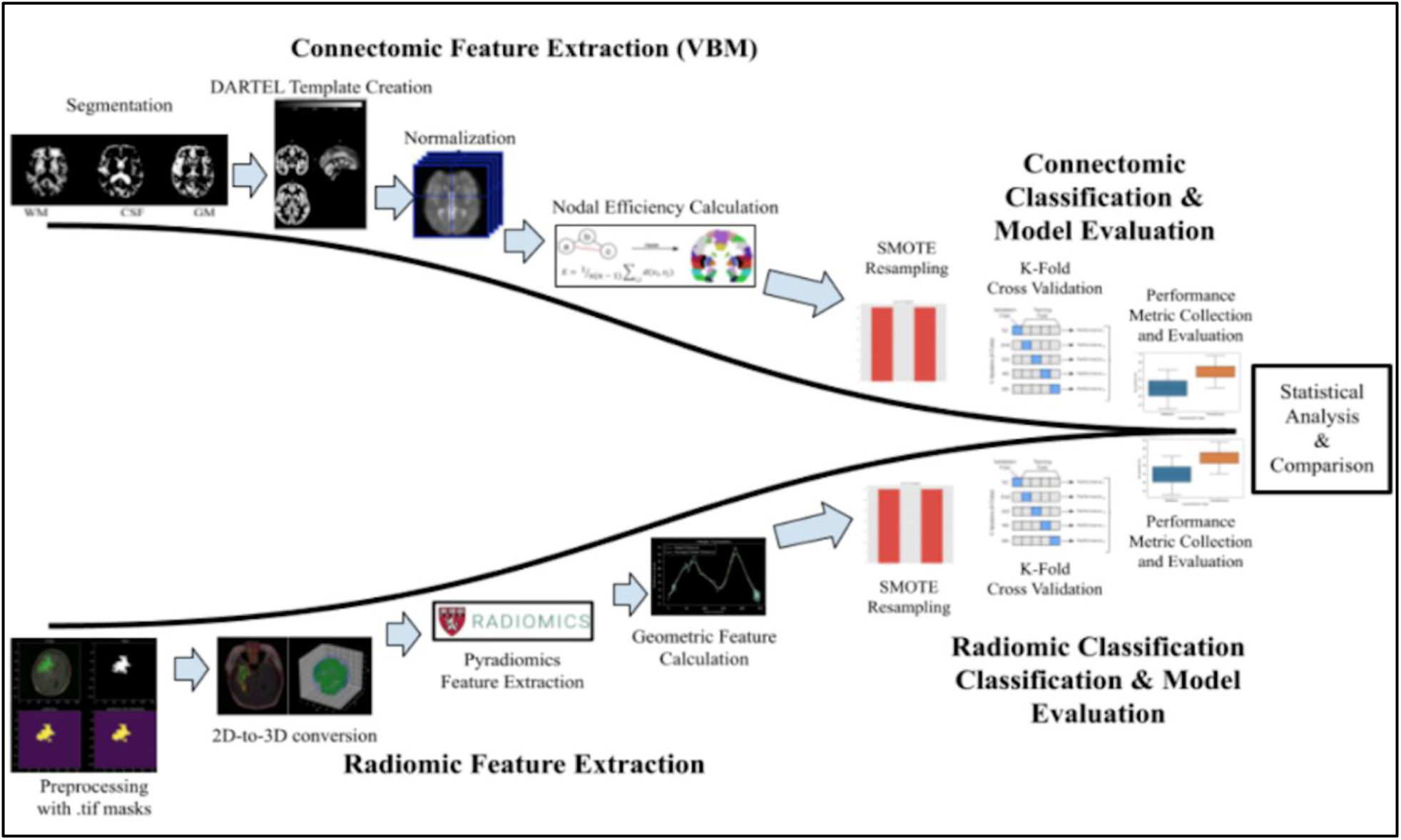
Workflow of experimental design and procedures ^14, 15, 16, 17^.

### Data and Patient Characteristics

In order to conduct this study, preoperative, precontrast imaging, corresponding ground truth IDH1 and 1p/19q status, and ground truth tumor masks were required. The necessary imaging data was sourced from The Cancer Imaging Archive (TCIA),^18^ which provided T1-weighted preoperative Magnetic Resonance Imaging (MRI) for 60 patients. Each patient folder contained between 30-88 slices. All genetic panel testing data came from The Cancer Genome Atlas (TCGA) and provided labels to document IDH1 and 1p/19q co-deletion status.^19^ Tumors with wild-type IDH were labeled as 0, and tumors with mutated IDH1 were labeled as 1. Similarly, 1p/19q co<u>-</u> deleted tumors were marked with a 1, and non-co<u>-</u>-deleted tumors were marked with a 0. In order to extract radiomic features relating to LGGs, region of interest (ROI) masks outlining the tumor were needed. These masks were sourced from a dataset of manual segmentations in the .tif format that were created and validated by radiologists.^20^ After all data had been acquired, patient ID was used to organize and reconcile among the three different sources. The dataset included patients with varying histological grades, tumor locations, gender, age at initial diagnosis, race, and ethnicity.

### Connectomic Feature Extraction

In order to extract the tumors’ connectomic features, two-dimensional patient images had to be converted into three-dimensional volumes using the tool, MRIcroGL. First, the 2D .dcm images were converted into the 3D .nii format. Next, these images were incorporated into a VBM pipeline^21^ in order to create gray matter-based structural connectomes. To do this, we employed the CAT12 toolbox^22^ which uses statistical parametric mapping (SPM) methods to perform skull-stripping, remove peripheral tissue, and segment gray matter, white matter, and cerebrospinal fluids (CSF). Smoothing and normalization techniques were applied to maintain imaging uniformity. Because of the high variance in brain size and shape, the DARTEL (Diffeomorphic Anatomical Registration using Exponentiated Lie algebra) toolbox^23^ was used to minimize subject disparity by aligning a sample-specific template to which patients’ images were fit. Using the AAL90 parcellation scheme, the gray matter files were distributed into 90 brain regions. The parcellated output gray matter files were then submitted to the BNets Toolbox,^24^ which used MATLAB scripts to calculate nodal efficiency for each region. Nodal efficiency was chosen as a connectome feature because this property has been shown to be affected in patients with diffuse glioma. ^14^ The network sizes (total number of nodes) and network degrees (total number of connections), and total brain volume were computed, resulting in a total of 93 connectomic features per patient.

### Radiomic Feature Extraction

We employed Python from the Pyradiomics^17^ package to extract 120 of the 123 total radiomic features. To do this, the 2D imaging was converted to 3D, this time using the JoinSeries conversion function which is part of the SimpleITK toolkit ^25^. Because there were 30-88 slices per patient, only tumors with the largest 3D volume for each patient were selected, as this would provide the best available representation of the LGG in 3D. Three other features, Bounding Ellipsoid Volume Ratio (BEVR), Angular Standard Deviation (ASD), and Margin Fluctuation (MF), were calculated geometrically through pixel manipulation due to their importance in classification based on other studies.^26^ BEVR was calculated using the Khachiyan Algorithm from an open source toolkit.^27^

### Tumor Classification

For both radiomics and connectomics, we created a binary classifier for IDH1 status and another for 1p/19q co<u>-</u>-deletion status. For the two connectomic classifiers, the input labels, X, for each patient were the 92 connectomic features that had been calculated in the previous steps. Similarly, the 108 radiomic features served as input labels for the two radiomic classifiers. For IDH1 classifiers, output labels (Y) were 0 for IDH1 wild type and 1 for mutated IDH1. The labels for 1p/19q co-deleted tumors were 1 and 0 for 1p/19q co-deleted and non-co-deleted, respectively. Within each classification model, eight different commonly used machine learning algorithms were implemented as follows: K-Nearest Neighbors (KNN), Logistic Regression, Naive Bayes, Support Vector Machine (SVM) with a radial basis function (RBF) kernel, Linear SVM, Decision Tree, Random Forest, and XGBoost. Because of the small sample size and data imbalance, several methods were employed to optimize performance. For all four classifiers, the best classification algorithm was selected for further improvement, refinement, hyperparameter tuning, and metric analysis. K-fold cross validation^28^, a resampling method commonly used on small datasets, was applied in order to improve classification. Using this method, the dataset is divided into *k* non-overlapping groups and each is used as a held-back test set to diversify model evaluation. In addition, the Synthetic Minority Oversampling Technique^29^ (SMOTE) was utilized to minimize the effects of data imbalance on performance. This technique up-samples the minority class through data augmentation. Afterwards, the model was trained and tested. For evaluation of model performance, the area under the receiver operating characteristic (AUROC), also known as the area under curve (AUC) score, and accuracy were tabulated for each classifier. In addition, feature importance values based on an importance metric were assigned to each input feature, both radiomic and connectomic, to form a ranked list of features based on their contribution to the classifier. The top radiomic and connectomic features, quantified by the importance metric, were used to perform a combined classification model. This combined classification followed the same procedure as the radiomic and connectomic models.

### Performance Evaluation Criteria

We employed both accuracy and AUC scoring in order to compare objectively the performance of our radiomic and connectomic techniques to predict IDH1 and 1p/19q co-deletion status. Accuracy quantifies how well a model classifies given a randomly selected test set, while the AUC determines the validity of a model’s decision threshold; that is, it communicates how effectively a model distinguishes between different classes based on the threshold. For the purposes of comparison, if one of the approaches (radiomic or connectomic) performed better than the other in both accuracy and AUC metrics, it was automatically considered a better classifier for that purpose. On the other hand, if neither classifier achieved both higher accuracy AND higher AUC score, resulting in draw, then statistical significance would be seen as a “tie-breaker”. In the case that there was a draw, and no statistically significant differences existed at all, model performance would be interpreted as equal.

### Data Analysis and Visualization

Box-and-whisker plots were drawn using the seaborn package in Python to represent accuracy distribution of classification over 10 runs. Additionally, the receiver operating characteristic (ROC) was drawn using matplotlib and its accompanying AUC score was plotted for the best AUC value attained by the model. Mean accuracy and AUC scores were calculated over 10 runs. To effectively compare the performance of the two forms of classification, two statistical tests, the Hanley-McNeil Test^30^ for AUC score comparison and the McNemar’s Test^31^ for accuracy comparison, were used to quantify statistical significance of differences in performance metrics using p-value. Finally, to visualize brain regions of interest based on importance to the binary classifiers, the BrainNet Viewer graphical user interface mapped relevant regions onto the AAL90 template.

## Results

As shown in Table 1, the mean age of patients 47.48 ± 12.69. There was the same number of males and females, 30, included in the patient dataset. In addition to these features, other important patient characteristics were summarized in Table 1: IDH1 and 1p/19q co<u>-</u> deletion status, Karnofsky Performance Status, or KPS, histologic grade, histological type, tumor laterality, and tumor location.

**Table 1.** Patient Characteristics.

|  |  |
| --- | --- |
|  | N = 60 |
| Age | Mean: 47.48 ± 12.69<br>Range: 22–70 |
| Sex | Male: 30 (50.00%) |
|  | Female: 30 (50.00%) |
| IDH1 Status | Wildtype: 20 (33.3%) |
|  | Mutated: 40 (66.6%) |
| 1p/19q Co-deletion Status | Co-deleted: 42 (70%) |
|  | Non-co-deleted: 18 (30%) |
| Karnofsky Performance Status (KPS) Score (N = 38) | 100: 12 (31.58%) |
|  | 90: 20 (52.63%) |
|  | 80: 4 (10.53%) |
|  | 70: 0 (0.00%) |
|  | 60: 0 (0.00%) |
|  | 50: 2 (5.26%) |
| Histologic Grade | Grade II: 25 (41.67%) |
|  | Grade III: 35 (58.33%) |
| Histological Type | Oligodendroglioma: 27 (45.00%) |
|  | Astrocytoma: 16 (26.67%) |
|  | Oligoastrocytoma: 17 (28.33%) |
| Tumor Laterality | Right: 34 (56.67%) |
|  | Left: 24 (40.00%) |
|  | Midline: 2 (3.33%) |
| Tumor Location | Frontal: 31 (51.67%) |
|  | Temporal: 22 (36.67%) |
|  | Parietal: 6 (10.00%) |
|  | Cerebellum: 1 (1.67%) |

Table 2 displays the performance metrics used to evaluate the two approaches. Each binary classifier was run 10 times, and the average AUC score and average accuracy were calculated. As can be seen, the radiomics method identified the 1p/19q co-deletion with a mean AUC score of 0.91 (scale: 0.00-1.00), a standard deviation (SD) of 0.11 and an 83.7% (2.8) mean accuracy. In contrast, the connectomic method identified the 1p/19q co-deletion with a mean AUC score of 0.70 (0.1) and was 86.7% (2.2) accurate. Radiomic classification of IDH1 yielded a mean AUC score of 0.83 (0.15) and a mean accuracy of 74.8% (1.4). while the connectomic classification of IDH1 status achieved a 0.85 mean AUC score (0.1) and a 77.0% (1.0) mean accuracy. The difference between the mean radiomic and connectomic AUC scores and accuracies were computed for both classifications. These differences as well as the statistical significance of the differences are recorded in the table as well.

**Table 2.**
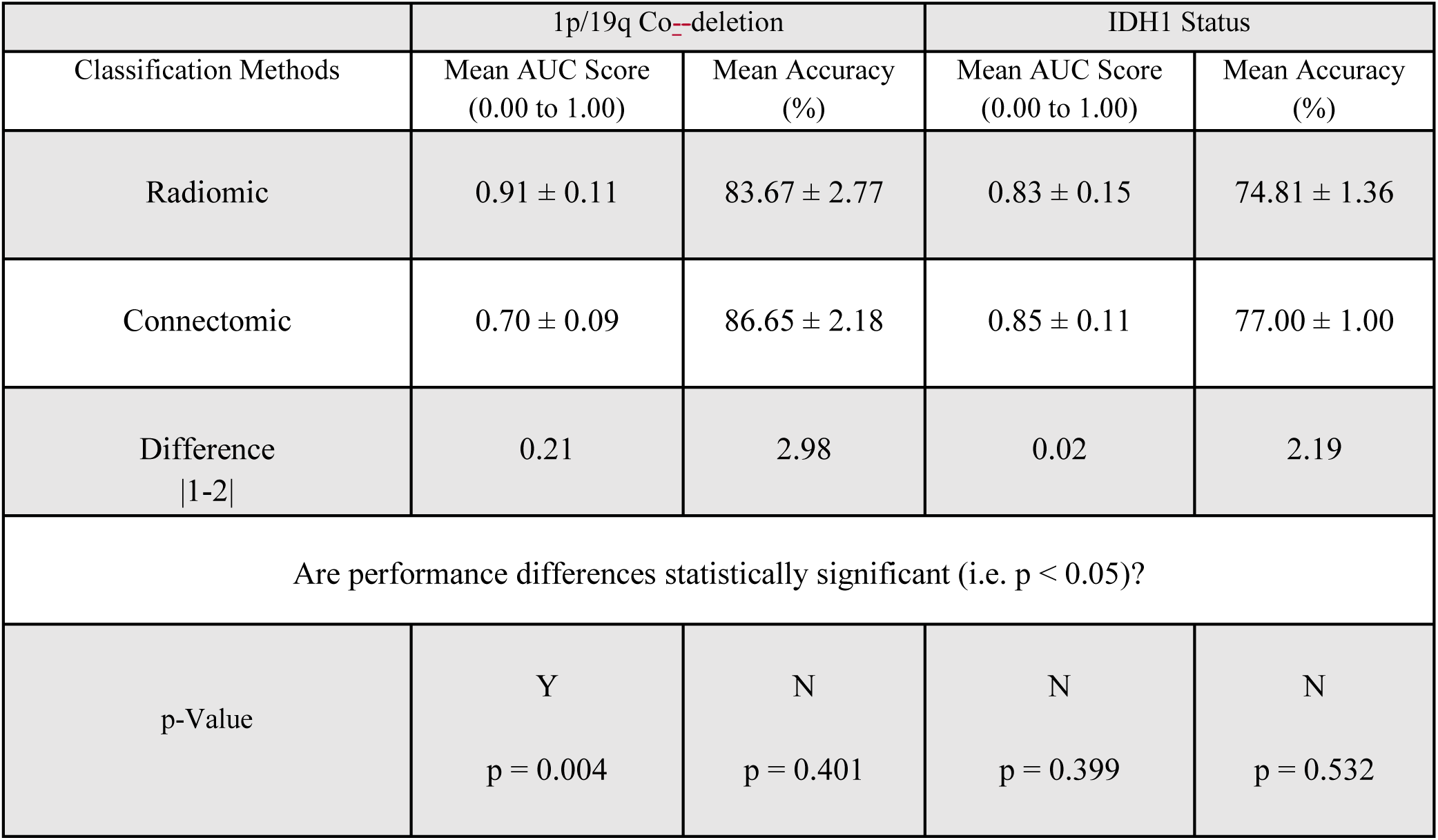
Comparison of mean classifier performance metrics for radiomic and connectomic approaches.

Overall, there was no significant difference between the mean accuracies of the radiomic and connectomic classifiers. With regards to mean AUC scores, the radiomic and connectomic classifiers for IDH1 status were comparable. The radiomic classifier for the 1p/19q co-deletion had a mean AUC score that was 0.21 greater than the connectomic classifier. This difference was the only one found to be statistically significant among performance metric comparisons.

Finding only minor differences in the performance of the two methods, we next tested whether or not the radiomic and connectomic approaches were complementary in which case, combining the methods might enhance predictive accuracy. This combined classification approach utilized the top five ranked radiomic and connectomic features for both 1p/19q co-deletion status as well as IDH1 status.

As shown in Table 3, this combined classification method achieved a mean AUC score of 0.93 with an SD of 0.02 and a mean accuracy of 86.0% (1.74) for identifying tumors with the 1p/19q co-deletion and a 0.85 (0.02) mean AUC score and a 78.0% (1.06) mean accuracy for identifying the IDH1 mutation.

**Table 3.** Mean classifier performance metrics for combined classification approach.

| Classification Methods | 1p/19q Co-deletion |  | IDH1 Status |  |
| --- | --- | --- | --- | --- |
|  | Mean AUC Score<br>(0.00 to 1.00) | Mean Accuracy<br>(%) | Mean AUC Score<br>(0.00 to 1.00) | Mean Accuracy<br>(%) |
| Combined Method | 0.93 ± 0.02 | 85.97 ± 1.74 | 0.85 ± 0.02 | 77.99 ± 1.06 |

Figures 4a and 4b display bar graphs comparing the mean AUC scores and mean accuracies of the combined (red), radiomic (orange), and connectomic (blue) classification models. Figure 4a shows that the mean AUC scores of the combined and radiomic classification models for 1p/19q co-deletion were 0.93 and 0.91, respectively. Both values were significantly larger (p = 0.004 for both) than the 0.70 mean AUC score of connectomic classification for 1p/19q co-deletion. Combined and connectomic models for IDH1 status classification both received mean AUC scores of 0.85, while the radiomic model’s mean AUC score was 0.83. (Fig 5a) No significant difference existed between mean AUC scores for IDH1 status classification. As shown by Figure 4b, combined, radiomic, and connectomic classification approaches for 1p/1q co-deletion had mean accuracies of 86.0%, 83.7%, and 86.7%, respectively. With respect to classification of IDH1 status, combined classification had higher mean accuracy (78.0%) than connectomic (77.0%) or radiomic classification (74.8%). (Fig 4b) For classification of 1p/19q co-deletion and IDH1 status, there were no significant differences in the mean accuracies of the three models.

**Figure 4a.**
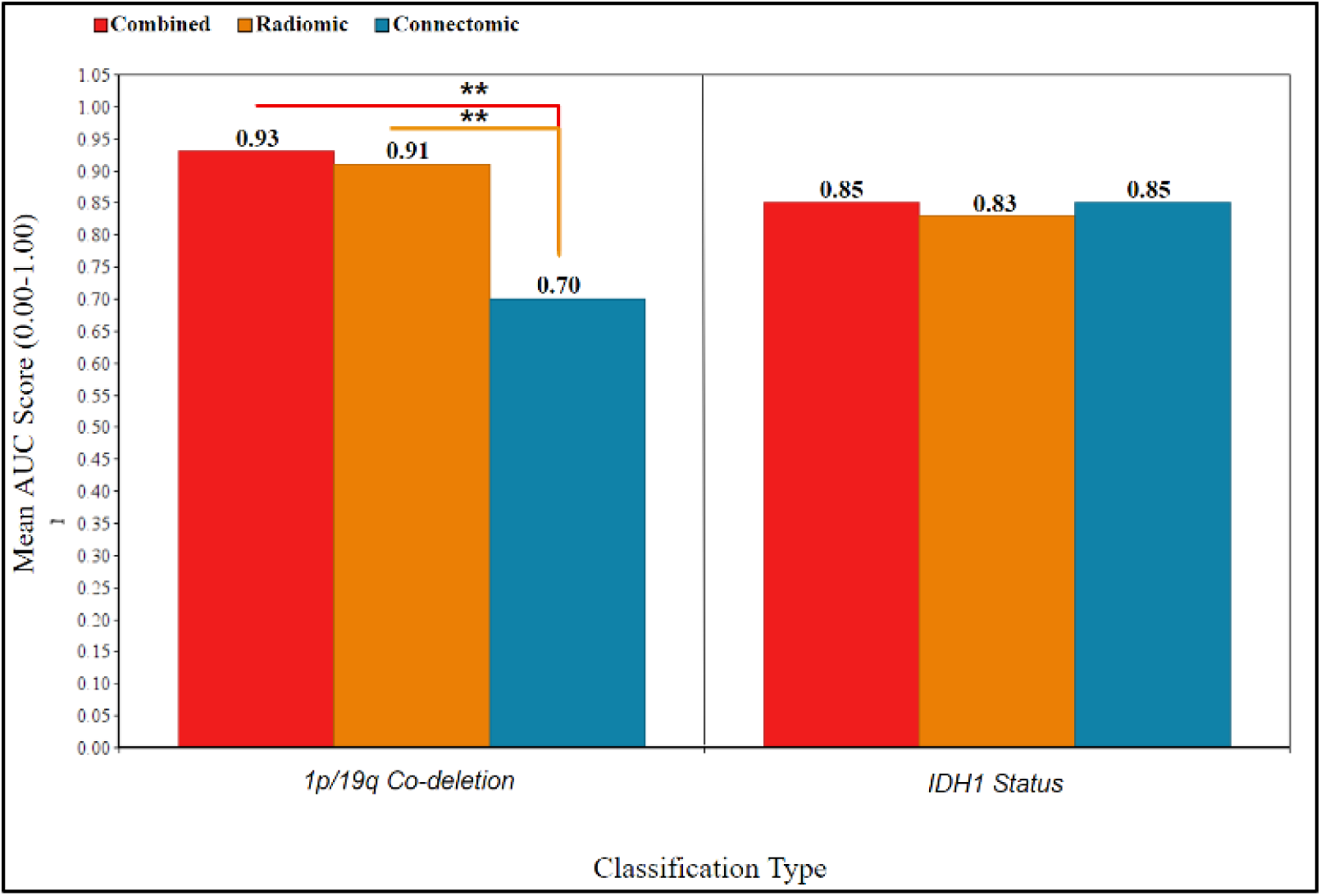
Comparison of mean AUC scores of combined, radiomic, and connectomic approaches to classification.

**Figure 4b.**
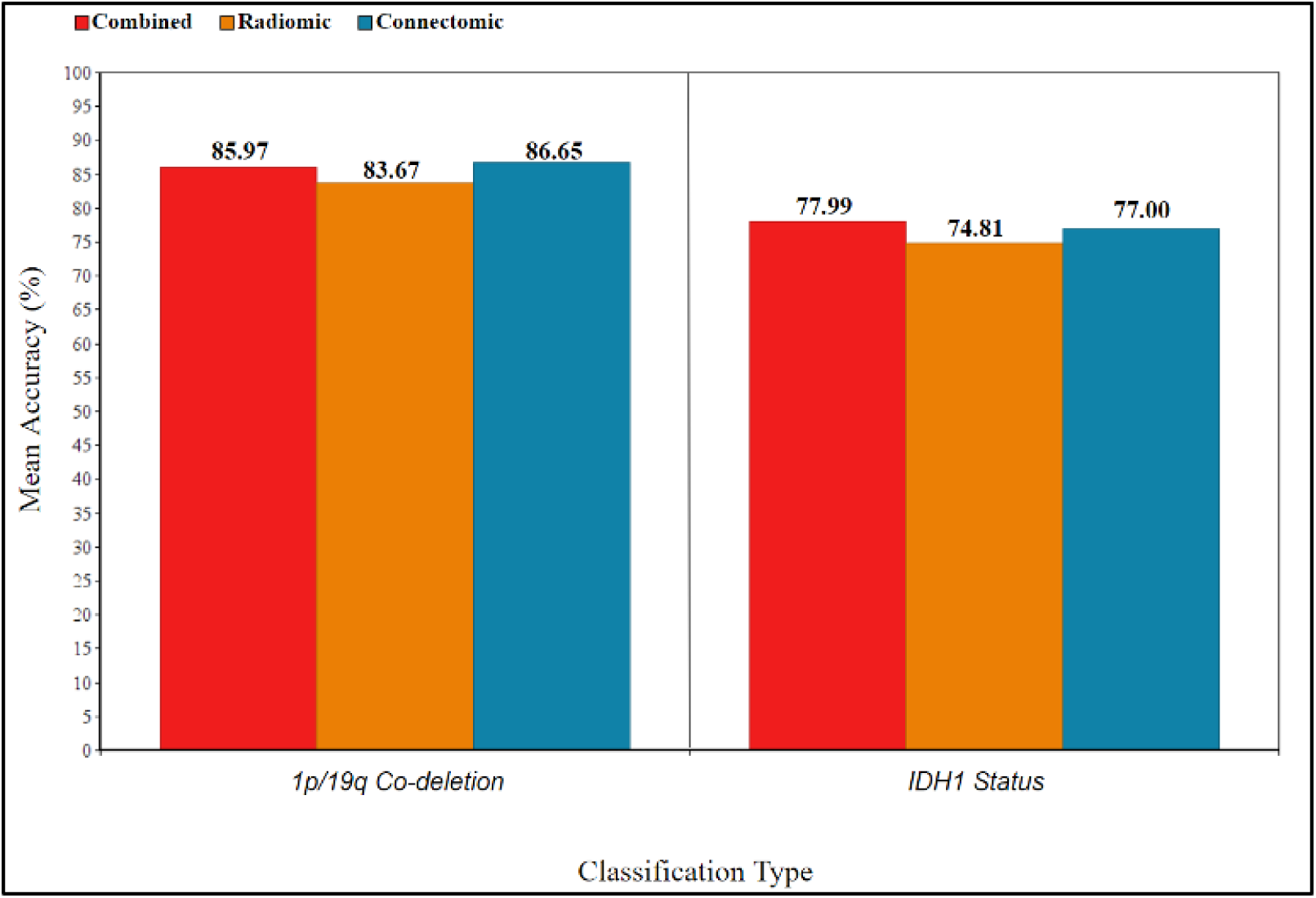
Comparison of mean accuracies of combined, radiomic, and connectomic approaches to classification.

**Figure 5a.**
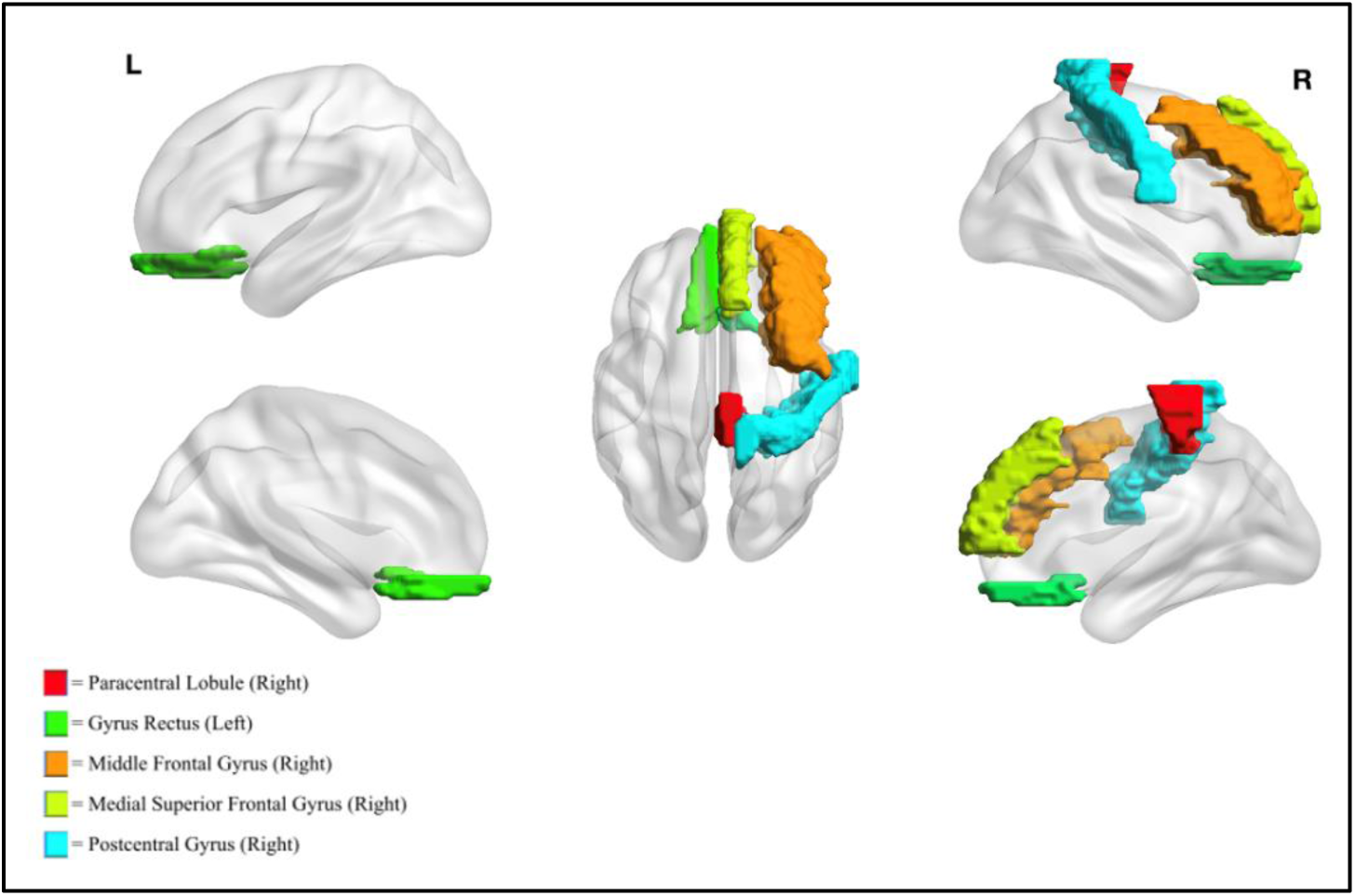
Visualization of top-five ranked brain regions based on IDH1 classification feature importance.

Finally, we identified those brain regions and radiomic features that had high relative importance in relation to classification of both IDH1 and 1p/19q co-deletion status. A feature importance metric was applied in all classifiers, and all of the brain regions and radiomic features in Tables 4a and 4b were found to have the most overall contribution to classification in terms of a feature importance metric.

**Table 4.**
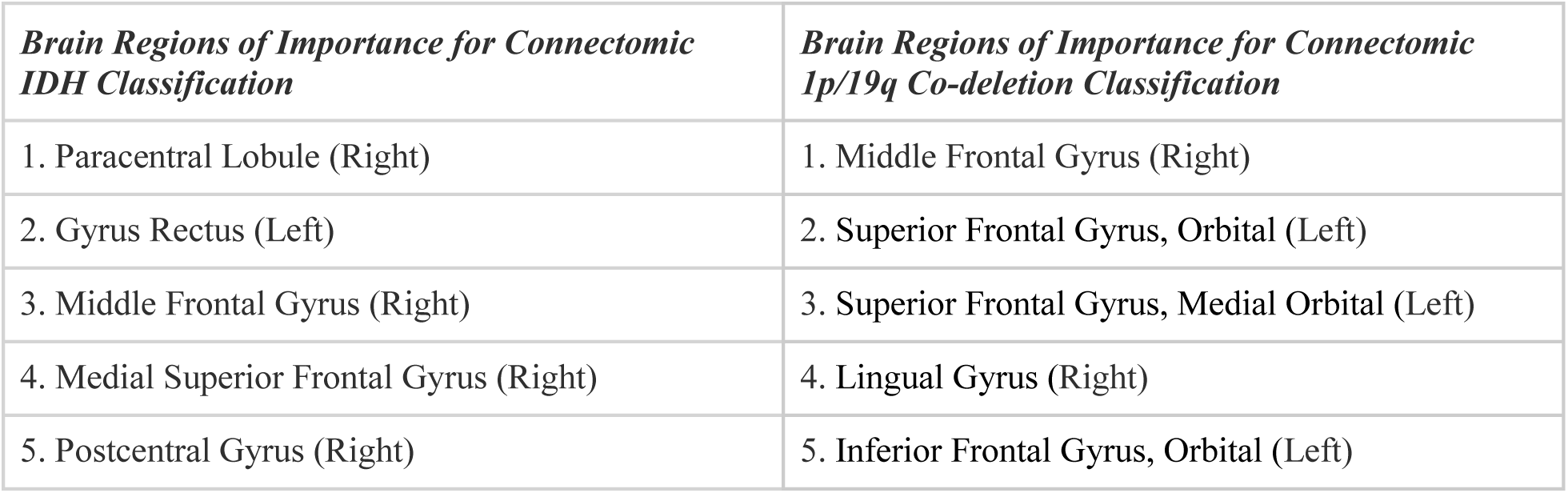
Top-five brain regions based on connectomic feature importance metric.

**Table 4b.** Top-five brain regions based on radiomic feature importance metric.

| *Brain Regions of Importance for Radiomic IDH Classification* | *Brain Regions of Importance for Radiomic 1p/19q Co-deletion* |
| --- | --- |
| 1. Gray Value Range | 1. Long Run High Gray-Level Emphasis |
| 2. Root Mean Squared (RMS) | 2. Inverse Difference Normalized (IDN) |
| 3. Uniformity | 3. Dependence Non-Uniformity Normalized |
| 4. Margin Fluctuation (MF) | 4. Zone Entropy (ZE) |
| 5. Dependence Entropy (DE) | 5. Maximal Correlation Coefficient (MCC) |

Figures 5a and 5b exhibit the brain regions that were considered highly important to the classification models according to this feature importance metric. The visualizations are brain regions mapped from the AAL90 parcellation scheme.

**Figure 5b.**
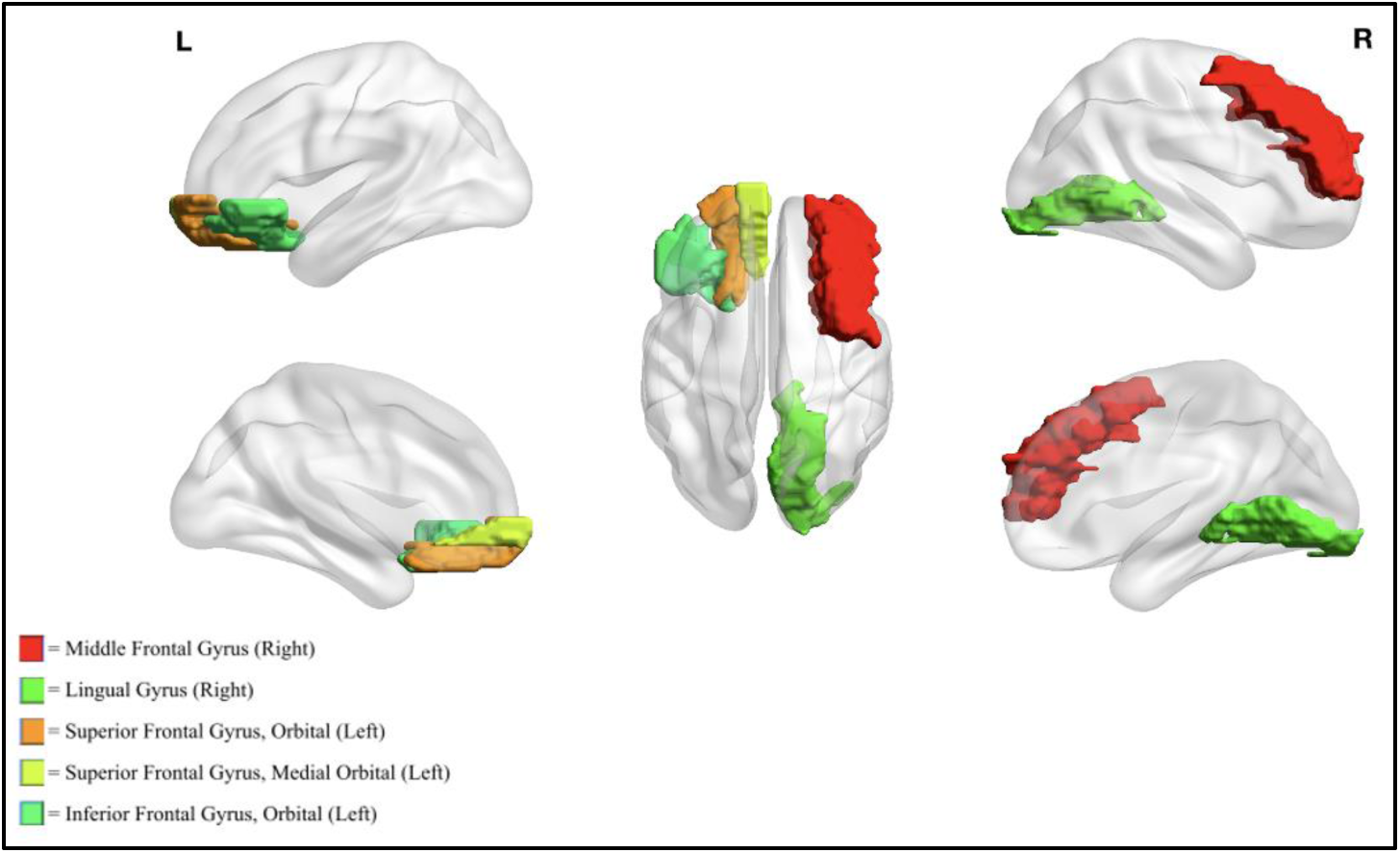
Visualization of top-five ranked brain regions based on 1p/19q co-deletion classification feature importance.

## Discussion

The purpose of this study was to compare the ability of radiomic and connectomic approaches to predict IDH1 and 1p/19q co-deletion status in a series of LGGs. Two objective metrics, mean accuracy and mean AUC score, were used to make this comparison, consistent with the prior literature. The main findings were as follows:

1. Both methods accurately predicted the IDH1 and 1p/19q co-deletion status of the tumors.
2. While the connectomic method yielded higher mean accuracies in both classification tasks, the differences between the two methods were not statistically significant.
3. The Radiomic approach had a signficantly higher mean AUC score than did the connectomic approach when predicting 1p/19q co-deletion status, the only statistically significant difference in performance between the two models.
4. Combining the approaches did not significantly improve the predictive performance of our model.

The classification models we developed for this study equaled or outperformed prior image-based models for classifying the 1p/19q co-deletion and IDH1 mutation status of gliomas. Zhou et al. extracted radiomic features from multi-modal MRIs to classify the 1p/19q co-deletion and IDH1 mutation status of 538 low- and high-grade gliomas. Their model was 78.2% accurate with an AUC score of 0.69 in the training set and 0.72 in the validation set, which was equal to or slightly lower than the mean accuracies and AUC scores of the classification models used in this study.^10^ Yu et al. performed radiomic classification of IDH1 with an 80% accuracy and 0.86 AUC score^11^; similar to the 77.0% mean accuracy and 0.85 mean AUC score achieved by IDH1 connectomic classification and the 78.0% mean accuracy and 0.85 mean AUC score generated by our combined classification model. One reason why both the radiomic and connectomic classification models in this study may have had slightly worse performance than the Yu et al. model is our smaller data set (60 vs 110 patients).

The results of this comparative analysis were also encouraging regarding the connectomic approach, which had only been attempted in one prior study and in that study only to predict IDH1 mutation satus.^14^ That study, performed by Kesler et al., yielded an 86% accuracy with a 0.94 AUC score.^14^ Again, the superior performance of their model as compared to ours may be attributed to a larger data set (234 patients). In addition, these authors employed 3T MRI, which has a higher tissue resolution than the 1.5T MRIs employed in our study.

In the current study, we found that the Random Forest classifier was the highest performing algorithm for connectomic and radiomic IDH1 classification as well as for radiomic 1p/19q classification. In contrast, the Linear SVM was the best method for connectomic 1p/19q classification. This supports the findings of previous studies ^9, 10, 11, 14^ and validates the effectiveness of the connectomic approach to classifying IDH1 and 1p/19q co-deletion status. Connectomic approaches allow for a new perspective to glioma classification, one that may reveal previously unobservable phenomena in the neurobiological effects of LGGs. Given the success of this and prior studies, we believe the connectomic approach merits further study, employing larger high resolution imaging sets and perhaps more robust algorithms. The only major detriment to real-world applications of the connectomic approach is the computational processing required to build connectomes. In this study, high performance computing with 48 CPUs still required run times of 4-6 hours per patient. Given the limited technology infrastructure of most hospitals, a great deal of work would need to be done to make this method more efficient. On the other hand, this may represent an opportunity for outside entities to provide tumor classification services to hospitals on a contractual basis.

Another contribution of this study is the discovery of specific radiomic and connectomic features that were important to classification (Tables 4a and 4b), something few prior studies have examined. Future studies may reveal how various brain regions and networks are affected by a tumor and impact functional outcomes or even survival after surgery. In addition, very few studies have identified features of importance for IDH1 and 1p/19q co-deletion status, so this study could serve as a precursor to experiments using radiomic and connectomic features to identify additional tumor biomarkers.

Perhaps the broadest implication of this study is that this research establishes a new framework under which IDH1 and 1p/19q co-deletion classification can operate. This is the first study that compared connectomic and radiomic methods using standard conditions and a uniform set of imaging data as controls. The finding that the two approaches were largely comparable leads us to recommend further exploration of collaborative, joint strategy to classification. A new direction of research, which incorporates both connectomic and radiomic features in classification models, such as the one conducted in this study, could be used to potentially outperform current efforts and become a viable alternative to current invasive measures.

Although this study provided many important findings, there were several limitations that may have affected accuracy, scope, and methodology. One of the major limitations was a lack of imaging data available that included all the requisite information needed to perform classification. Because imaging, tumor masks, and labels for 1p/19q co-deletion and IDH1 mutation needed to be cross-referenced, data availability was a major hurdle. Although many steps were taken to minimize the effects of our small sample size and data imbalance, there is no doubt that larger sample sets yield more reliable information. Furthermore, because this experiment was performed using the exact same patient samples, some data points had to be omitted because they did not include one of the required items. These factors may have affected the scope of the study as well as the metrics that defined model performance. Another limitation that may have affected the methodology arose from the unavailability of studies performing connectomic classification. Because there was just one published study on classification of IDH1 using a connectomic approach, developing an accurate experimental design was a challenge. With almost no prior research on the topic, there was only one existing methodology to reference and use as a starting point for this study. Finally, the importance of our findings of feature importance metrics of brain regions for prediction 1p/19q co-deletion and IDH1 mutation may be limited in scope. This is because feature importance can only provide information on how useful a predictor is within model constraints—these metrics cannot supply any insights into the significance of these brain regions in classification of IDH1 status and 1p/19q co-deletion unless further investigation is undertaken.

Although the results of this study were very encouraging, data unavailability could have affected their quality. One recommendation for future research is to replicate the methods described in this study using a larger, more comprehensive dataset. Furthermore, model inputs could be restricted to include only radiomic and connectomic features ranked highly for importance, according to Tables 4a and 4b. Another consideration would be including glioblastoma multiforme (GBM), a grade IV astrocytoma whose prognosis and treatment response is, like LGG, dictated by IDH1 status and 1p/19q. One course of action building from this and other studies could be to replicate the pipeline model created by Lu et al.^9^ using findings from this study. This convoluted pipeline operates in three steps: classification to discriminate between LGG and GBM, IDH1 classification, and 1p/19q classification. Lu et al. performed this model using only radiomic classification, however, a consolidated approach in which IDH1 classification is done through connectomics and 1p/19q co-deletion is classified through radiomics could lead to higher model performance.

We have no Conflict of Interest

